# Self-Driven Jamming in Growing Microbial Populations

**DOI:** 10.1101/052480

**Authors:** Morgan Delarue, Jörn Hartung, Carl Schreck, Pawel Gniewek, Lucy Hu, Stephan Herminghaus, Oskar Hallatschek

## Abstract

In natural settings, microbes tend to grow in dense populations [1–4] where they need to push against their surroundings to accommodate space for new cells. The associated contact forces play a critical role in a variety of population-level processes, including biofilm formation [5–7], the colonization of porous media [8, 9], and the invasion of biological tissues [10–12]. Although mechanical forces have been characterized at the single cell level [13–16], it remains elusive how collective pushing forces result from the combination of single cell forces. Here, we reveal a collective mechanism of confinement, which we call self-driven jamming, that promotes the build-up of large mechanical pressures in microbial populations. Microfluidic experiments on budding yeast populations in space-limited environments show that self-driven jamming arises from the gradual formation and sudden collapse of force chains driven by microbial proliferation, extending the framework of driven granular matter [17–20]. The resulting contact pressures can become large enough to slow down cell growth by delaying the cell cycle in the G1 phase and to strain or even destroy the microenvironment through crack propagation. Our results suggest that self-driven jamming and build-up of large mechanical pressures is a natural tendency of microbes growing in confined spaces, contributing to microbial pathogenesis and biofouling [21–26].

---

The simulataneous measurement of the physiology and mechanics of microbes is enabled by a microfluidic bioreactor [27–30] that we have designed to culture microbes under tightly controlled chemical and mechanical conditions. The setup, shown in Fig. 1A, is optimized for budding yeast (*S. cerevisiae*). We use this device to measure mechanical forces generated by partially-confined growing populations and the impact of those forces on both the population itself and its micro-environment.

**Figure 1.**
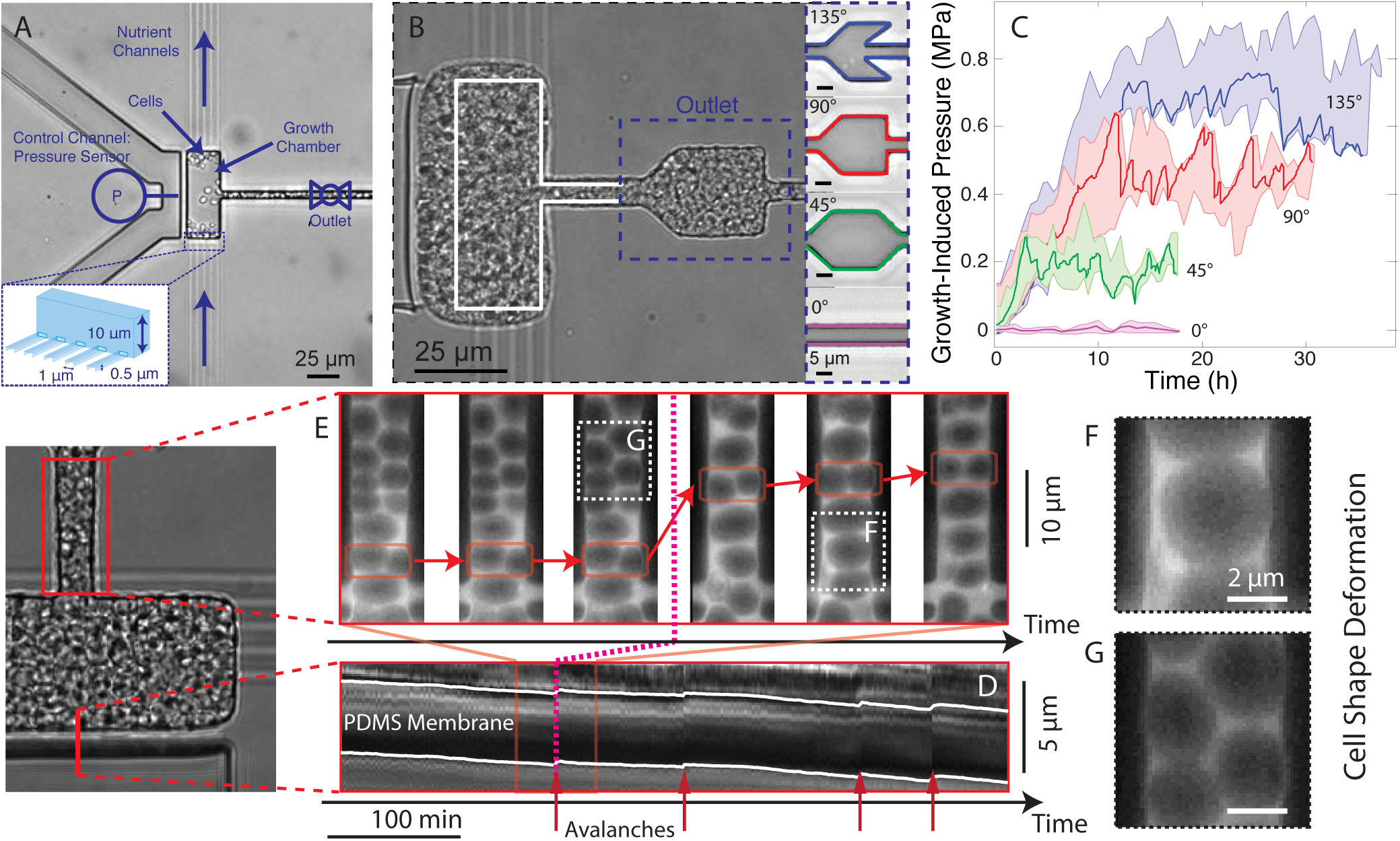
Self-driven jamming of microbes enables collective pressure build-up in microfluidic environments. (**A**) Budding yeast cells are grown in a growth chamber threaded by narrow nutrient channels (inset). (**B**) The jamming of excess microbes produced by proliferation in the device leads to a partial confinement of the population and a gradual build-up of a contact pressure of up to 0.65 ± 0.1 MPa (in the shown experiment), which strongly deforms the device (white line represents the undeformed layout). The steady-state pressure generated in a given device depends on the geometry of the outlets (**B**, right), which effectively act as leaky one-way valves. The resulting time-dependent pressure curves are shown in (**C**) for different outlets. The pressure measurements were enabled by an automatic feedback system that actively controls the deformation of a thin membrane separating the growth chamber and a control channel (see **A** and Supplementary Text). The bold curves correspond to one realization of the experiment, which is characterized by large pressure fluctuations due to gradual jamming and sudden unjamming. The shaded region represents the envelope of the replicates: all replicates are binned together, and within each bin, the minimum and the maximum define the shading. The cellular flows exhibits collective features known from physics of jamming in granular media: The outflow of cells is not steady but consists of periods of stasis, accompanied by pressure-build up, and sudden cell avalanches and pressure drops. This can be seen in time lapse movies ( **here**) as well as Kimographs: (**D**) shows the random zig-zag motion of the chamber membrane and (**E**) shows the flow through the outlet before, during and after an avalanche with one snapshot every 20 minutes. Note that, depending on the local stresses, cells can assume shapes from nearly spherical (**F**, low stress) to nearly polyhedral (**G**, high stress).

At the beginning of each experiment, we trap a single yeast cell in the growth chamber of the device, which can hold up to about 100 cells. The cells are fed by a continuous flow of culture medium, provided by a narrow set of channels that are impassable for cells.

While cells first proliferate exponentially as in liquid culture, growth dynamics change dramatically once the chamber is filled. At high density, cells move in a stop-and-go manner and increasingly push against the chamber walls. The population develops a contact pressure^*^ that increases over time until it reaches steady state, subject to large fluctuations. Depending on the geometry of the device (Fig. 1B and C), the mean steady-state pressure can reach up to 1 MPa. This pressure is larger than the ≈ 0.2 MPa turgor pressure measured in budding yeast (stationary phase [31]) and much larger than the ≈ 1 mPa needed for the cells to overcome viscous friction (supplementary text).

Both the intermittent flow and pressure build-up are counterintuitive because the outlet channel is wide enough for cells to pass. In principle, excess cells could flow like a liquid out of the chamber. Time lapse movies (**here**) reveal that blockages in the device stabilize the cell packing and prevent flow. Cells proliferate until a sudden avalanche flushes them through the outlet (Fig. 1D and E). Another jamming event occurs, and the process repeats. These dynamics generate characteristic slow pressure increases followed by sudden pressure drops (Fig. 1C).

Jamming, intermittency and avalanches are familiar aspects of flowing sand, grains or even jelly beans [24]. To test whether the interplay of growth, collective rearrangement, and outflow of cells from the chamber can be explained by the mechanics of granular materials, we set up coarse-grained computer simulations with cells represented as elastic particles that grow exponentially and reproduce by budding. In our simulations, cells move via frictionless overdamped dynamics with repulsive contact interactions between neighbors.

Our simulations indeed reproduce the intermittent dynamics observed in the experiments (Fig. 2A–C). We find that the pressure drops are roughly exponentially-distributed for both experiments and simulations (Fig. 2D) for *P* > 〈*P*〉, consistent with hopper flows [32].

**Figure 2.**
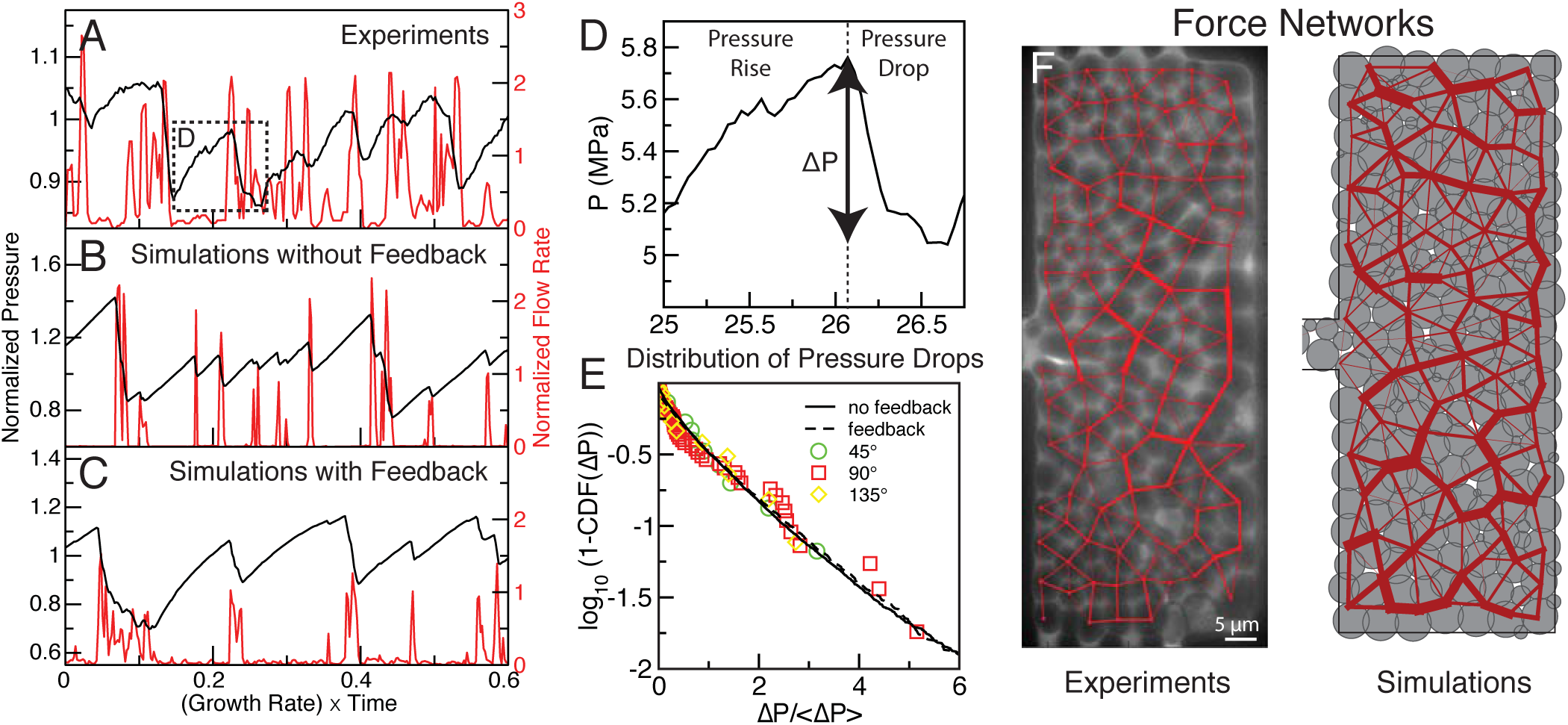
Pressure fluctuations and intermittent flows of partially confined budding yeast populations can be reproduced in simulations of proliferating elastic particles. (**A**) Experimental pressure time series are characterized by periods of gradual pressure build-up and sudden pressure drops. (**B**) Simulations show that such time series are the generic outcome of jammed elastic particles proliferating in confined spaces. (**C**) A feedback of pressure onto growth, reported in Fig. 3C below, further improves our simulations. The gradual pressure increases prior to avalanche events show diminishing return similar to the experimental time series in (**A**). Pressure drops during avalanche events (**D**) are nearly exponentially distributed for drops larger than the mean pressure drop, 〈Δ*P*〉, in both experiments (**E**: symbols) and coarse-grained simulations (**E**: lines). We can estimate inter-cell contact forces in our experiments by measuring the area of contact between two cells through image analysis. (**F**) The resulting network of contact forces in packings of budding yeast cells shows a heterogeneous distribution of mechanical stresses (pressure on the membrane: 0.5 MPa). (**G**) Force networks obtained from simulations of exponentially growing budding cells. In both (**F**) and (**G**), large forces are clustered into chain-like structures. A movie illustrating the dynamics of force networks in our experiments can be seen **here**, and a coarse-grained simulation movie can be seen **here**. For our simulations, we used box and outlet sizes that match the microfludic chamber and parameterized the over-damped dynamics using the experimental flow rate and pressure fluctuation data (supplementary text).

**Figure 3.**
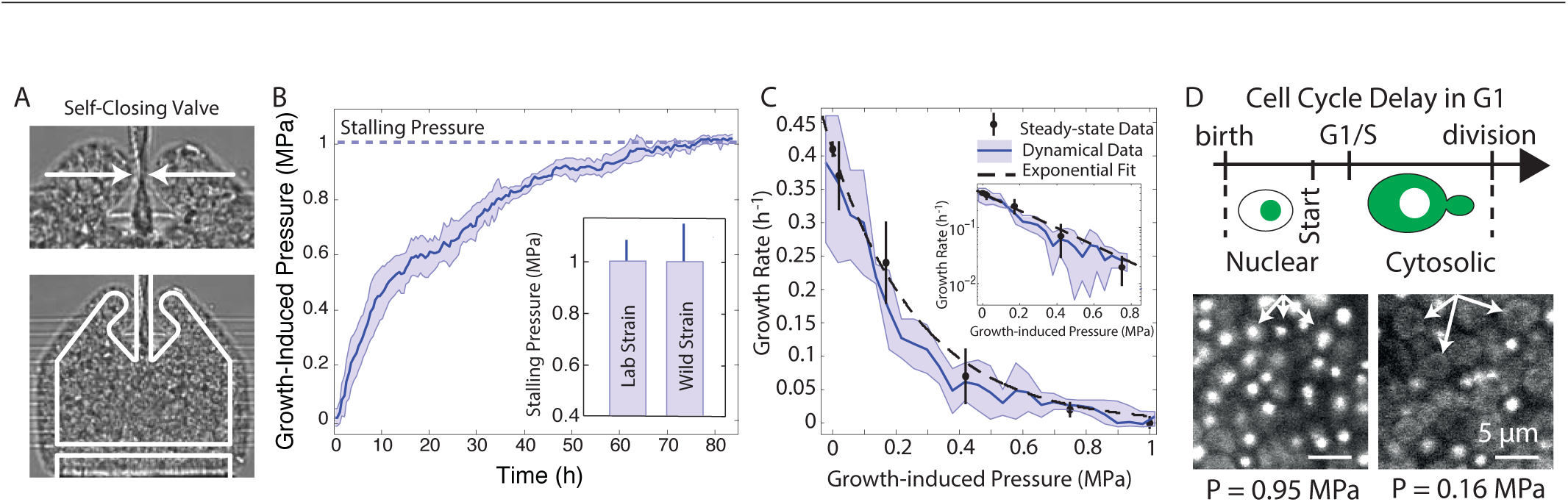
Pressure-induced slow down of growth. (**A**) Budding yeast populations can be fully confined using a “self-closing” device that takes advantage of the contact pressure developed by the population to close the inlet/outlet channel. The cells are fed through narrow nutrient channels, as in 1**A**. The layout of the undeformed device is shown in white. (**B**) The time-dependent pressure curve in the self-closing devices shows a diminishing return: The rate of increase of the growth-induced pressure in the fully confined region gradually slows until it stops at the stalling pressure of 1±0.1 MPa (5 replicates). Inset: stalling pressure measured for the lab strain and the wild strain. (**C**) Growth rate as a function of growth-induced pressure, estimated in two ways (supplementary text): The black points represent net growth rates determined from the cell flow out of our leaky devices in the steady-state (black points; ≥ 5 replicates). The continous blue line, on the other hand, has been inferred from the deminishing return in the dynamical data of (**B**) under a quasi-steady state assumption (supplementary text; shading indicates ± SD). The dashed curves represents an exponential fit to the steady-state data (k = 0.41 (h^−1^)exp(-P/0.28 (MPa))). (**D**) Cells under pressure spent a larger fraction of time in G1 phase, as inferred from the fraction of cells presenting a nuclear Whi5 signal.

Highly intermittent cell flows might reflect spatially heterogeneous mechanical stresses, a hallmark of driven granular materials [17–20]. Assuming that cell shape deformation is indicative of the forces between cells, we developed a non-invasive method to infer these forces (Fig. 2F, supplementary text, and fig. S1). Using this approach, we analyzed microscopy images to determine stress distributions of crowded populations. Both *S. cerevisiae* experiments and our coarse-grained simulations exhibit disordered cell packings that are stabilized by heterogeneous force networks (Fig. 2F and G). Stress is highly localized along branching “force chains” [17, 18] while adjacent “spectator cells” [33] experience very little mechanical stress.

We find that jamming-induced contact forces can become so large that they feed back on the cell physiology. Indeed, a feedback on both cell shape and the dynamics of cell growth is evident in experiments where we place two devices of different steady state pressures next to one another, as seen in the time lapse movie (**here**). To quantify the feedback on growth, we estimate the net growth rate, which is the difference between birth and death rate, in our microfluidic bioreactors by measuring mean cell outflow rate at steady state (supplementary text). We find that the growth rate decays roughly exponentially with pressure until growth is undetectable at a stalling pressure of about 1 MPa (Fig. 3C). The stalling pressure, or homeostatic pressure [34], is obtained by using a special device with a “self-closing valve”, in which yeast populations fully confine themselves by the pressure they build up, as seen in Fig. 3A. In this device, the rate of pressure increase gradually decays with pressure until saturation (Fig. 3B). This diminishing return is due to smaller growth rates at higher pressures, and serves as another, dynamical measure for the feedback between contact pressure and growth rate.

Control experiments supported by finite element simulations show that cells are well-fed and viable even at the highest densities suggesting a mechanobiological origin for the reduced growth rates (supplementary text and figs. S3 and S4). As a first step to uncover the mechanistic basis for the feedback, we have found that contact pressure acts to slow down the cell cycle in the G1 phase (Fig. 3D). Specifically the fraction of cells in G1, indicated by subcellular localization of the protein Whi5, increases with decreasing growth rate (fig. S5). This result consistent with a recent study showing a cell cycle arrest in G1 in compressed mammalian cells [35].

Perhaps the most salient consequence of growth-induced pressure is cell shape deformations. While budding yeast cells grown in the absence of mechanical stresses are nearly spherical, we observe that they morph into convex polyhedra as the population pressure becomes growth-limiting (Fig. 1F and G). Close to the stalling pressure, the packing resembles the structure of a dry foam [36], consisting of cells with only flat faces and sharp edges in between, shown in Fig. 2F. The pressure-induced cell shape deformation can be best visualized at the interface between coverslip and cell population: the cell-coverslip contact area increases as the growth-induced pressure increases (Fig. S6). Our simulations further suggest that the cell turgor pressure in the experiments may increase as a function of the growth-induced pressure.

Most microbial cells are sticky [37, 38]. Indeed, while our lab strains of budding yeast have been domesticated to become non-sticky, wild strains can have strong, velcro-like intercellular fiber connections [39]. We find that while these sticky strains develop a very similar maximal pressure as the lab strains do (Fig. 3B), they develop growth-induced pressures under much weaker confinement (Fig. 4A). Our coarse-grained simulations further suggest that attraction between cells can lead to a build up of pressure much larger than expected under a null model of a liquid droplet with surface tension (Fig. 4C and D).

**Figure 4.**
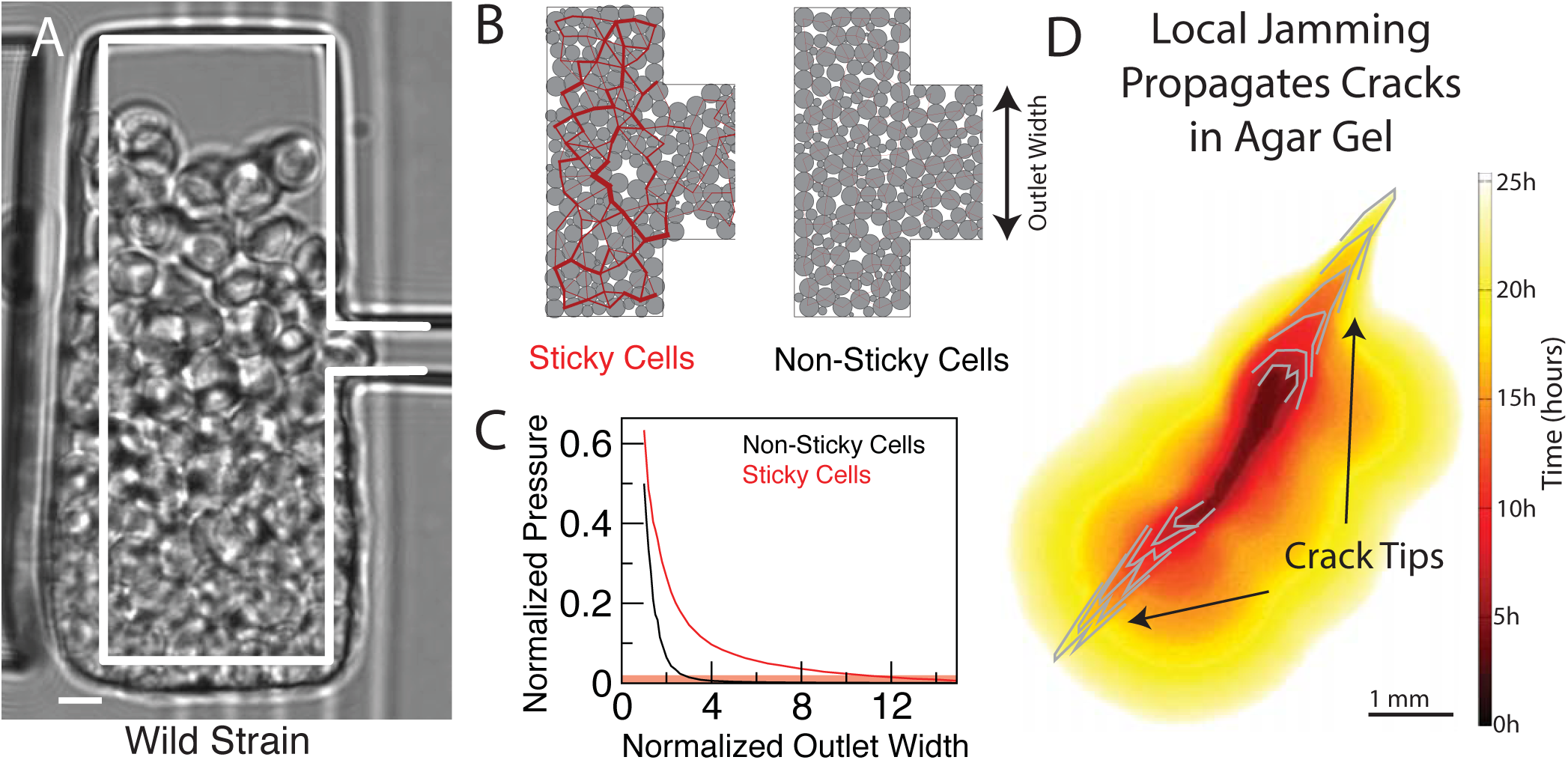
Self-driven jamming is promoted by stickiness and can remodel the microenvironment. (**A**) Wild strains of yeast stick together via strong velcro-like connections between cells [39], This stabilizes the spherical growth of the population against shear stresses. (**B, C**) Simulations show that even weak attractive forces between cells can strongly promote jamming. (**B**) Packing of slightly sticky cells (right, supplementary text) exhibit a force network with pronounced force chains in contrast to the non-sticky case for the shown device. (**C**) The increase in growth-induced pressure (steady-state) with stickiness is much larger than expected from the continuum limit (red base line) over a broad range of outlet sizes (supplementary text). (**D**) Gradual propagation of agar gel cracks by growing populations of budding yeast (lab strain). Cells grow out of a pre-exisiting agar crack and, at the same time, propagate the crack tips inside the agar. A time-lapse movie of the crack propagation is available **here**.

Bacteria and fungi have the ability to colonize a wide range of porous media, including tiny cavities barely larger than their cell size [3, 4]. Our work suggests that self-driven jamming of growing microbes can emerge in these microenvironments as it does in our microfluidic devices if chemical resources are sufficiently abundant. The mechanism underlying self-driven jamming is cell proliferation, thus extending the notion of jamming, which usually result from external sources of driving, such as shear, compression, or gravity [17–20].

The resulting growth-induced forces endow biofilms with the potential to remodel, or even destroy, their micro-environment. This could aid microbes in penetrating the soft tissues of host organisms [10–12], or to invade soil, where most microbes grow in pores of several micro-meter in diameters [3, 4]. At this length scale, it is possible that the growth-induced pressures measured here contribute to straining of even stiff materials. Indeed, when we grow budding yeast populations inside agar gels, we observe the formation and propagation of cracks (Fig. 4D, fig. S8 and time lapse movie **here**). Thus, just like jamming of granular media can threaten the mechanical integrity of their confinements, which can lead to the bursting of grain silos [32, 40], it could also be an important mechanical aspect of host invasion [10–12] and biofouling [21].

## Acknowledgments

We would like to thank Jasper Rine, Jeremy Thorner and Liam Holt for helpful discussions. Research reported in this publication was supported by the National Institute Of General Medical Sciences of the National Institutes of Health under Award Number R01GM115851, by a Simons Investigator award from the Simons Foundation (O.H.) and by the German Research Foundation (DFG) in the framework of the SFB 937/A15. The content is solely the responsibility of the authors and does not necessarily represent the official views of the National Institutes of Health.

## Author Contributions

O.H. designed and supervised the study. M.D., J.H., S.H. and O.H. designed the microfluidic experiments, M.D. and J.H. developed the software and performed experiments. M.D., J.H., and L.H. fabricated devices. M.D. and L.H. performed Comsol simulations, P.G. implemented and performed the mass-spring simulations, and C.S. implemented and performed the coarsegrained simulations. M.D., J.H., C.S., P.G. and O.H. interpreted the data and wrote the manuscript.

## Supplementary Material

**Yeast strains and growth conditions:** *S. cerevisiae* cells (haploid strain of mating type a, containing a GFP fused to Whi5 in a S288C background, obtained from J. Thorner laboratory at UC Berkeley) and wild undomesticated cells (BR-103F strain from the Palkova lab at Charles University in Czech Republic) are cultured in complete synthetic medium (CSM, 20 *g*/L glucose) at 30° C. Cells in exponential phase are loaded in the device.

**Preparation of the micro-fluidic bioreactor (“Mechano-chemostat”):** The master consists of 2 layers of different heights, prepared using classical soft-lithography protocole, as in [1] for instance. The first layer is prepared using SU 2000.5 negative photoresist (0.5 *μ*m height), and the second using SU 2010 (10 *μ*m height). Polydimethylsiloxane (PDMS, Sylgard 184, Dow Corning, USA) is mixed with the curing agent (ratio 1:10 in mass), poured onto the master, and cured overnight in a 60° C oven. It is bound to no1 thickness glass slides through an oxygen plasma generated by a reactive ion etcher (RIE) machine (P_02_ = 200 mTorr, exposure time = 20 sec). Prior to loading the device, the surface is treated with Pluronics 127 (VWR, USA) as in [2] to decrease any non-specific adhesion that could result in cell-PDMS friction. This ensures that the build-up of pressure is not due to adhesion or friction of the cells with the PDMS, but due to compressive forces between cells and the PDMS chamber.

**Measuring the growth-induced pressure:** We use the 4*μ*m thick membrane separating the growth chamber and the control channel, where the hydrostatic pressure can be imposed, to measure the contact pressure generated by the population. As observed in Fig. 1, cell growth deforms the membrane. We monitor the position of the membrane every 30 seconds, and change the hydrostatic pressure in the control channel to keep the membrane position constant, at all time. The membrane is thus in a mechanical equilibrium, and the imposed hydrostatic pressure in the control channel is balancing the growth-induced pressure in the growth chamber. The pressure can also be measured through the deformation of the membrane, finite element simulations (Comsol) showing that the deformation is proportional to the pressure. However, this requires a calibration of the Young’s modulus of the PDMS device. We measure an average PDMS elasticity of 2 MPa.

**Visualizing cell deformations and the contact area between cells and coverslip:** FITC-conjugated Dextran (3kDa, Invitrogen) is added to the culture medium, at a concentration of 0.1 mg/mL. Since Dextran is not internalized by single yeast cells [3], it stains the extracellular space, and enables the imaging of cell deformation. The contact between cells and the coverslip (fig.) is imaged by reflectometry. Briefly, we shine a 635nm laser on the sample through a pinhole closed to a minimum, to obtain an optical slice of 0.3 *μ*m. The reflected light is collected without filter, so that local changes in optical density can be measured at the level of the glass slide. Typical images of cell deformations are shown in the main Letter (Fig. 1), and images obtained by reflectometry are shown in fig. S6A.

**Measuring the steady-state and instantaneous growth rate:** Each outlet design shown in Fig. 1B leads to a different steady-state pressure, and a different steady-state cell outflow rate. We measure the cell outflow rate, *j*_cell_, using a custom-made particle image velocimetry algorithm (Matlab), and define the growth rate in the chamber as *k* = *j*_cell_/V_ch_, where V_ch_ is the volume of the growth chamber.

Alternatively, we can estimate the instantaneous growth rate from the pressure vs. time relationship measured for the self-closing device. Since the cells are fully trapped in the growth chamber, the time-derivative of the pressure is directly proportional to the growth rate. The proportionality depends on the packing fraction of the cells (*ϕ*) and on the volume of the chamber (*V*).

We define the instantaneous growth rate *γ* of the cells by

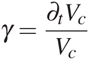

where *V*_*c*_ is the volume occupied by the cells. By definition, the packing fraction is the fraction of volume occupied by cells divided by the volume of the chamber:

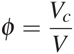

Hence,

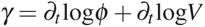

Now we assume that, at any time, the packing fraction and the chamber volume only depend on the pressure: *V*(*t*) = *V* (*P*(*t*)) and *ϕ*(*t*) = *ϕ* (*P*(*t*)). This quasi-steady state assumption is acceptable only if the cells can adapt their growth rates sufficiently fast to the current pressure curve or, conversely, that the pressure changes sufficiently slowly.

This enables us to rewrite the growth rate:

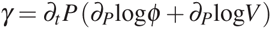

In order to plot the growth rate *γ* as a function of growth-induced pressure, we need three pieces of information: the time-derivative of the pressure, the packing fraction, and the pressure-dependency of the volume of the growth chamber. The packing fraction is measured using exclusion fluorescence technique (see Supplementary figure S2A and S2C), and the dependency on pressure of the volume of the chamber is calculated through finite element simulations (Comsol) (Supplementary Fig S2B).

We observe (fig. S2D) that the growth-rate vs pressure relationship obtained in this way is in good agreement with the more direct steady-state measurements. This justifies our steady-state assumptions and suggests that the feedback on growth should act as fast or faster than the typical division time.

**Inferring force maps:** The interface area between cells in contact is used to estimate the contact force between the cells. To this end, we have modeled the mechanical response of budding yeast cells in the simplest possible way by assuming that a cell responds to contact forces like a pressurized elastic shell, as illustrated in Fig. 2F. The force between cells in contact is then given by *F* = *PA* ∝ *pl*^2^, where *A* is the area of contact, *P* is the cell turgor pressure, and *l* is the projection of the contact surface onto the measurement plane. This takes into account the effects of turgor pressure and the near-inextensibility of the cell wall, but assumes that these effects dominate over elastic energies due to bending of the cell wall or cytoskeleton (the turgor pressure of ≈ 0.2 MPa [4] is nearly two orders of magnitude larger than the elastic moduli of cytoskeletal networks). Single-cell studies [5–8] have indeed found that compressed *S. cerevisiae* cells exert forces proportional to the area of contact, in agreement with a model that incorporates only internal pressure and cell wall stretching even for large deformations. We further validate our approach by performing simulations of deformable cells composed of spring networks, which show similar deformations as *S. cerevisiae* cells at corresponding pressures. The simulations are described in the next paragraph and in fig. 1.

**Description of Mass-Spring simulation:** The mechanics of a budding yeast cell is primarily controlled by the mechanics of the cell wall and the turgor pressure [8]. In our “mass-spring” (MS) simulations, the cell wall is represented as a spherical meshwork of springs obtained from surface triangulation. The neighbor vertices, separated by a distance **R**, are held together via Hookean spring interactions: **F**(**R**) = -*k*_MS_**R**(1 - *R*/*R*_0_), where *k*_MS_ is a spring constant. The overlap between two non-bonded vertices (or a vertex and a box wall) is modeled by Hertzian repulsive force: 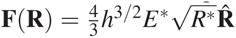 where: 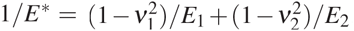;*E*_1_ and *E*_2_ are Young’s moduli; *v*_1_ and *v*_2_ Pois son’s ratios; 1/*R*\* = 1/*R*_1_ + 1/*R*_2_;*R*_1_ and *R*_2_ radii of the vertices, here representing cell wall thickness *t*, (*R*_2_ = ∞ for the box wall); and *h* = *R*_1_ + *R*_2_-*R*. The force due to the cell volume-dependent turgor pressure Π(V_cell_) on vertex *i* is calculated as 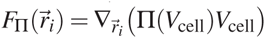, where 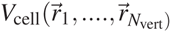 is a function of the *N*_vert_ vertices triangulating the cell surface and the volume change for vertex *i* is calculated using tetrahedral volume defined by vertex *i* and it’s neighboring 3 vertices. The equations of motion of over-damped dynamics have been solved using Heun’s method (explicit second-order Runge-Kutta method). The compression simulations have been performed by successive reduction of the size of the box with the rate 0.01-0.1*μ*m*s*^−1^. The parameters used in the simulations are: cell wall (box wall) Young’s modulus E=150 MPa (200 MPa); Poisson’s ratio v = 1/2; turgor pressure Π = 1.0 MPa (unless stated otherwise), t = 0.1*μ*m, initial radius of the cell *R*_0_ = 2.5*μ*m. The Hookean spring constants are taken to be the same and related to the Young’s modulus by the following equation: 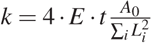, where E is Young’s modulus, *t* is the cell wall thickness, *A*_0_ is the cell area, and *L*_*i*_, is the relaxed length of the *i*^*th*^ spring [9].

**Dependence of contact surface area on pressure:** We measure the cell contact area at the interface between the coverslip and cell population, and compare it to in-house simulations. Reflectometry reveals that the average fraction of the coverslip that is in contact with cells increases as the population pressure increases, shown in fig. S6A. We find that the growth-induced pressure increases super-linearly with surface coverage, contradicting a pressurized-shell null model and suggesting the the yeast cell turgor pressure increases with growth induced pressure (fig. S6B).

**Coarse-grained simulations of proliferating elastic particles:** In our 2D coarse-grained simulations, illustrated in Supplementary fig. S9, cells are modeled as two frictionless rigidly-attached spherical lobes [10] (mother and bud) that grow exponentially at rate *γ*_*i*_ by bud expansion (Eq. 1), move according to over-damped dynamics with mobility *μ* (Eqs. 2 and 3), and interact via repulsive spring forces with elastic modulus *k* (Eq. 4)

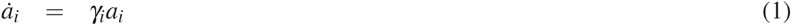

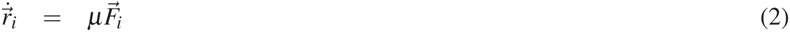

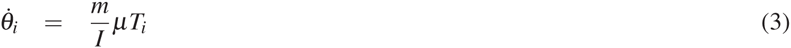

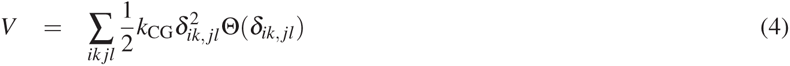

where 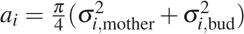 is the cell area, σ_*i,mother*_ (σ_*i,bud*_) is the diameter of the mother (bud), 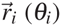 is the cell position (orientation), 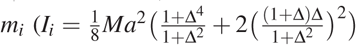 where Δ*i* = σ_*i,bud*_/σ_*i,mother*_) is the cell mass (inertia), *V* is the total potential energy, 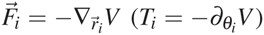 is the force (torque) on cell *i*, and 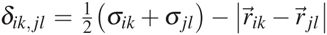 is the overlap between lobes *k* of cell *i* and *l* of cell *j*, and Θ is the Heaviside Step function. This method is similar to studies performed with growing spherocylinders [11, 52]. For simulations with attraction, we extend the potential in Eq. 4 beyond its repulsive core to have an attractive range of width Δ[13, 14]

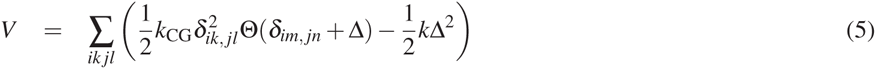

In this model, the mother lobe has the same size σ_*i*,mother_ = σ for all cells. Equations of motions are integrated using a 3^rd^ order Gear Predictor-Corrector algorithm. Growth progresses while σ_*i*,bud_ < σ and culminates in division where the daughter cells retain the orientation of cell *i*.

Cells grow in a rectangular box of dimensions *L*_*x*_ × *L*_*y*_ with an outlet of width *a*. For the simulations in this paper, we used *L*_*x*_ =6σ, *L*_*y*_ = 6σ, and *a* = 1.4σ to match experiments. Cells interact with the wall with the same force as other particles, 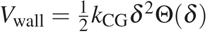, where *δ* is the overlap between the cell and wall.

Without pressure feedback, 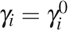 where 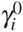 is chosen from a uniform distribution of width 20% around a mean growth rate *γ*. With pressure feedback, the growth rate depends on pressure as 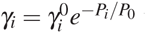 where *P*_*i*_ = is the pressure on particle *i*.

The free parameters in this model are an effective friction coefficient 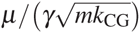 and a characteristic pressure feedback scale *P*_0_/*k*. In Fig. 3 of the main text, we use parameters that best matches the experimental pressure fluctuations in the case of intermittent flow where the pressure slowly builds and then suddenly drops during avalanches. We choose values of 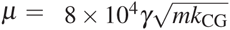 and 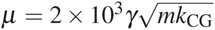 for simulations with (Fig. 2B) and without (Fig. 2C) feedback that best capture the ratio of pressure increase (*Ṗ*_↑_) and drop (*Ṗ*_↓_) rates in the case as shown in fig. S10. To obtain an experimentally-motivated value of feedback pressure *P*_0_ (Fig. 2C), we used a value of *P*_0_ that yields the same ratio of 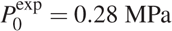 (Fig. 3C) to 〈P〉^exp^ = 0.7 MPa (135° data in Fig. 1C), 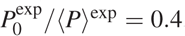. Coarse-grained simulations without feedback yield 〈P〉^sim^ = 0.19*k*_CG_, giving 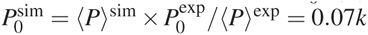

**Estimation of pressure due to viscous friction:** Here we estimate the pressure arising from friction between cells in the outlet and the surrounding medium. In a chamber of dimensions *L*_*x*_ × *L*_*y*_ with an outlet of dimensions width×length= a×d, the chamber holds *N*_*c*_ ≈ *L*_*x*_*L*_*y*_h/σ^3^ cells and the outlet holds *N*_o_ ≈ *adh*/σ^3^ cells, where σ is a typical cell diameter and *h* is the height of the device. Assuming that the height *h* of the system and the width of the outlet *a* are both *a* = *h* = σ, so that *N*_*c*_ ≈ *L*_*x*_*L*_*y*_/σ^2^ and *N*_o_ ≈ *ad*/σ^2^. If the cells in the outlet are pushed out at velocity *v*, the total frictional force they feel is *F* = *fvN*_o_, where *ƒ* is a friction coefficient per cell, and therefore the pressure at the outlet is

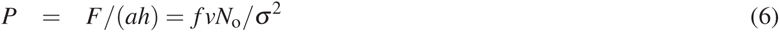

Standard viscous friction of a sphere in a liquid yields *ƒ* = 6*πη*σ/2 We further estimate the flow velocity by *v* = *N*_*c*_σ*k* where *k* is the growth rate for cells in the chamber, assuming that cells in the outlet are not growing. This gives:

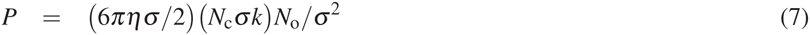

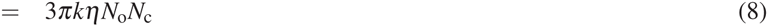

Using *η* = 10^−3^Pa s, *k* ≈ 0.4h^−1^ ≈ 10^−4^s^−1^, *N*_*c*_ ≈ 100, and *N*_o_ ≈ 10, we get

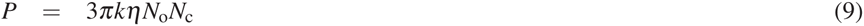

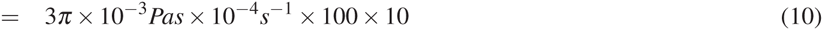

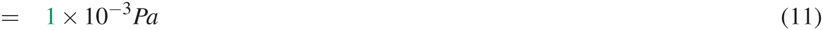

Thus, viscous friction gives a negligible contribution to the pressure generated in the outlet, which is in the MPa range.

Conversely, we can use the above estimate to define an effective viscosity of the cell packing of 1 MPa s needed to achieve a pressure of 1 MPa. This effective viscosity is much larger than has been measured for mammalian cells [15].

## Supplementary Figures

**Figure S1.**
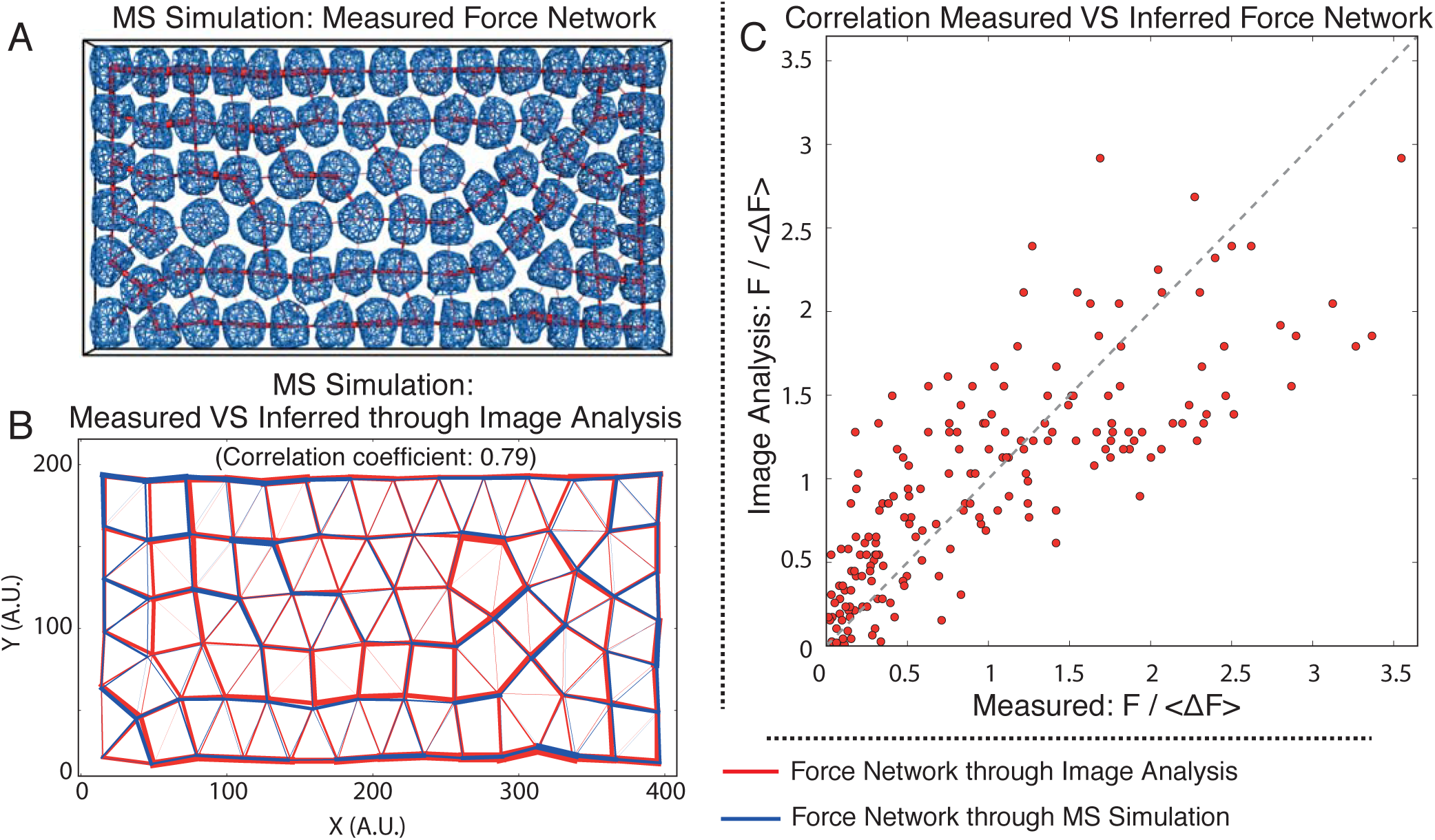
Testing our indirect force-inference method on simulated packings. In the main text, we reported mechanical stress distributions in packings of yeast cells that we have inferred from the observed cell shape deformations. Our force-inference method consists of the following steps: First, we used a custom Matlab image analysis code to process the time-lapse movies that we have obtained with the fluorescence exclusion method (fig. S2). Each cell, is identified with a watershed algorithm and manually refined if necessary. For each identified cell, the contour is defined as a set of spline functions. These splines are further used to calculate the length *l* of the contact line between each pair of cells. As a first order approximation, we estimate the contact area as *A* ∝ *l*^2^, and we assume that the contact force is proportional to the contact area *F* ∝ *A* (Materials and Methods: See Inferring force maps). Here, we test our force-inference method on packings generated by our mass-spring simulation. To this end, we compare the inferred force network with the actual force network in the simulations. **A**. 80 cells of the same size (*R*_0_ = 1.5*μ*m), turgor pressure (Π = 1.0 MPa), and E=100 MPa are randomly distributed and compressed in a slab geometry. The cells are depicted as a semi-transparent blue meshwork, confined by the rigid box. The contact forces are evaluated numerically and are represented as the red lines between neighbor cells. The thickness of the lines corresponds to the magnitude of the contact forces. **B**. The final snapshot from the simulation is processed with the in-house Matlab code for image analysis, and contact forces have been inferred. The numerical (in blue) and image analysis (in red) force networks are superimposed on top of each other for visual comparison. The correlation coefficient calculated for these two sets of contact forces is 0.79. **C**. Scatter plot of each contact force in **B**. Forces have been scaled by the average value. Measured are the forces obtained from the mass-spring simulations, and compared against the one obtained from the image analysis procedure.

**Figure S2.**
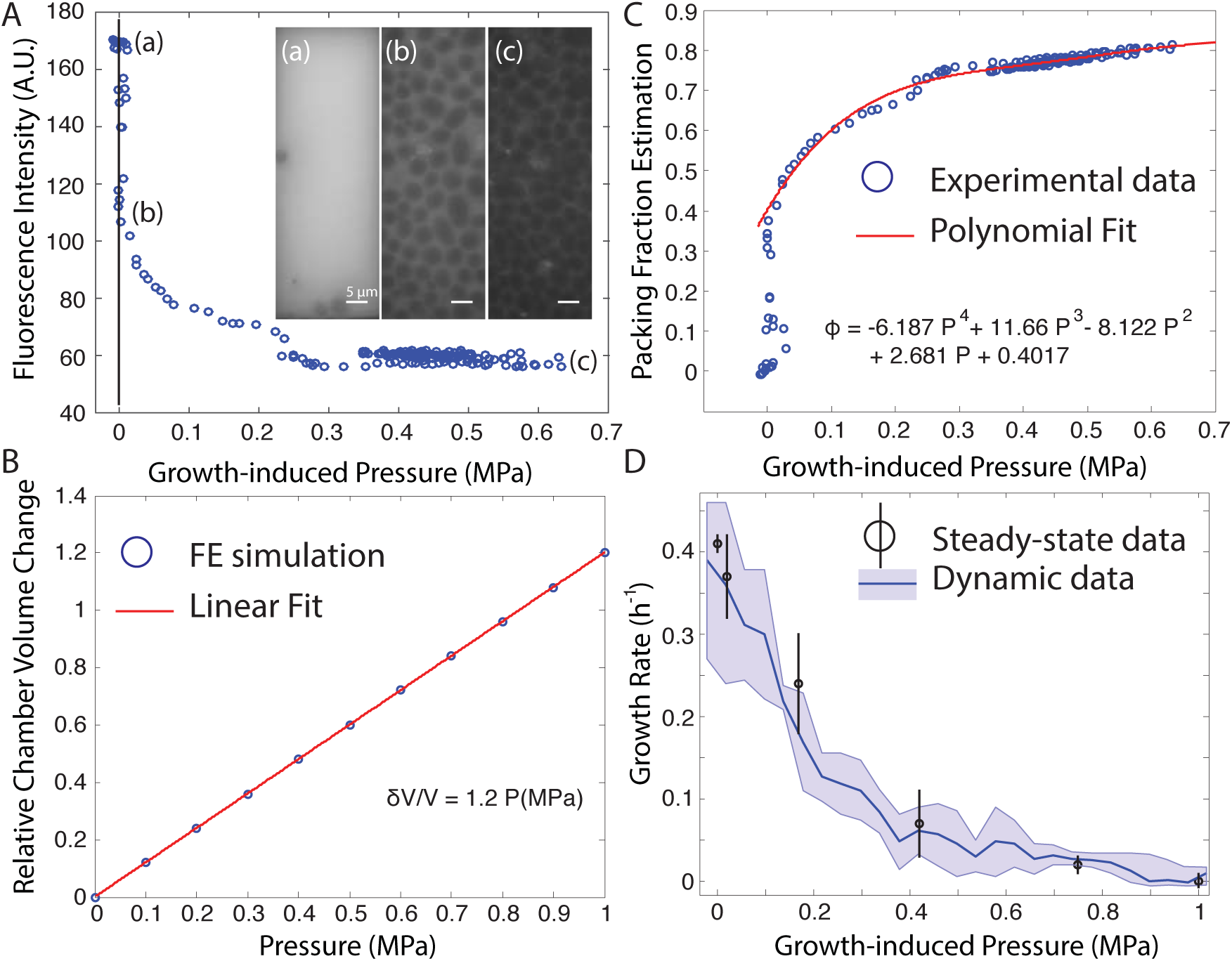
Inferring the instantaneous growth rate as a function of pressure using the pressure curve obtained from the self-closing valve. **A**. A fluorescent dye, FITC-conjugated Dextran, added to the medium allows us to label the space between the cells. FITC-conjugated Dextran does not penetrate inside the cells, such that its fluorescence is excluded from a cell. As a consequence, as the cells are filling the chamber, the fluorescence intensity is, in first order, proportional to the void in between cells, like in the fluorescent exclusion method [16]. Denoting *ϕ* the packing fraction, and *V* the volume of the chamber, we assume that the intensity *I* of fluorophore is *I* ∝ (1 – *ϕ*)*V*. **B**. We use finite element simulations (Comsol) to estimate the change in volume of the growth chamber as a function of the pressure. We define the PDMS as a hyperelastic material as in [17], with an estimated Young’s modulus E = 2MPa. We find that the change in volume is to good approximation linear in the pressure. **C**. We use the excluded fluorescence, as well as the finite element simulation, to estimate the cell packing fraction, *ϕ*, as a function of the growth-induced pressure. We observe that the growth-induced pressure starts to rise in the chamber for a packing fraction of about 0.4. We fit the resulting relationship by a forth order polynomial to obtain a continuously differentiable function. **D**. We use the values extracted from B and C to calculate the instantaneous volumetric growth rate *γ*, using a quasi-steady state assumption as described in the supplementary text. The dark blue line corresponds to the values calculated from the mean pressure, and the envelope corresponds to the values calculated from the envelope of the pressure curve. Note that the inferred continuous relationship between growth rate and contact pressure is in good agreement with the steady-state data obtained independently, from outflow rates in our leaky devices.

**Figure S3.**
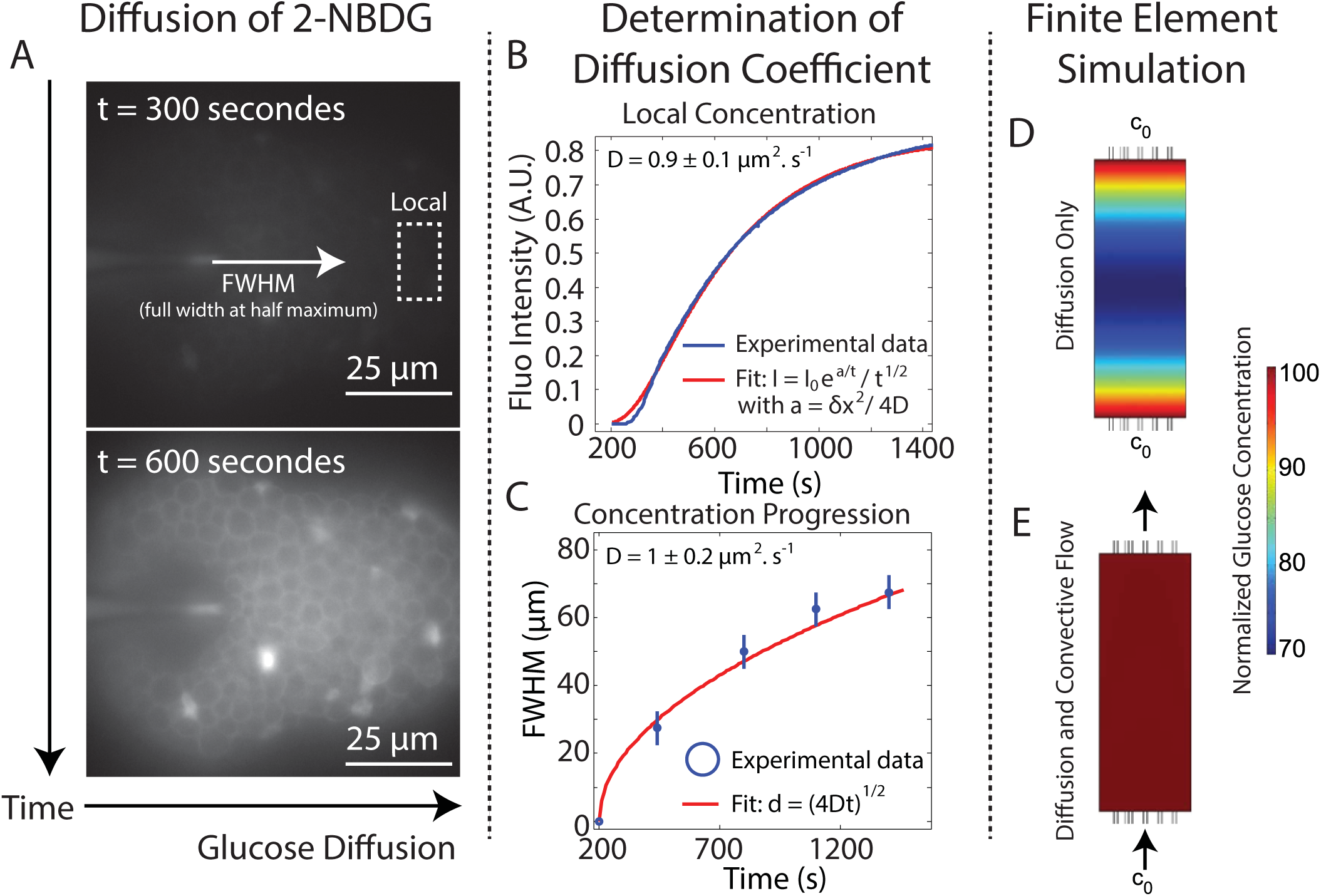
The reduction of growth rate is not due to glucose depletion in the growth chamber. To estimate whether glucose depletion could account for the observed reduction of growth rate, we assume that cells would locally consume glucose at the maximum rate. We consider two cases: either glucose merely diffuses inside the growth chamber, or it is also advected by the imposed nutrient flow. In both cases, we find that the reduction of glucose concentration in the chamber is not enough to stall cell growth. **A**. We first measure the diffusion of 2-NDBG, a fluorescent glucose analog molecule. Here, we observe at the beginning of the experiment that there is almost no glucose in the self closing valve, and that it progressively diffuses in the chamber. Notice the foam-like packing of the cells, which results from the growth-induced pressure nearly balancing the turgor pressure. **B - C**. We measure the diffusion constant of the glucose analog in 2 different ways. We measure either the local concentration at a fixed position in the chamber (**B**) or the full width at half maximum (FWHM) as a function of time (**C**). Fitting of a simple diffusion model agrees well with the experimental data, and enables us to extract values for the diffusion constant of the glucose analog (see figure). **D - E**. The biomass yield of *S. cerevisiae* cells is 0.45 × g_cells_/g_glucose_ [18]. With a minimum doubling time of 2 hours, this yields a glucose consumption rate of 2.2 × 10^7^ molecules/s. We simulate glucose consumption in the fully packed growth chamber using finite element simulations (Comsol) and the measured glucose diffusion constant extracted in **B** and **C**. We consider two cases: either there is only consumption and diffusion (**D**) or consumption, diffusion and convection (**E**). We find that in the case where there is only diffusion, the glucose concentration drops at about 70% of its boundary value *c*_0_, which is about 14 g/L, and still above the concentration where depletion of glucose affects growth [19]. In a finite element simulation set-up where we impose a convective flow of 0.2 nL/s, we observe that there is no glucose gradient in the growth chamber. We conclude that the observed reduction of growth rate in figure 3C is not an effect of glucose depletion in the growth chamber.

**Figure S4.**
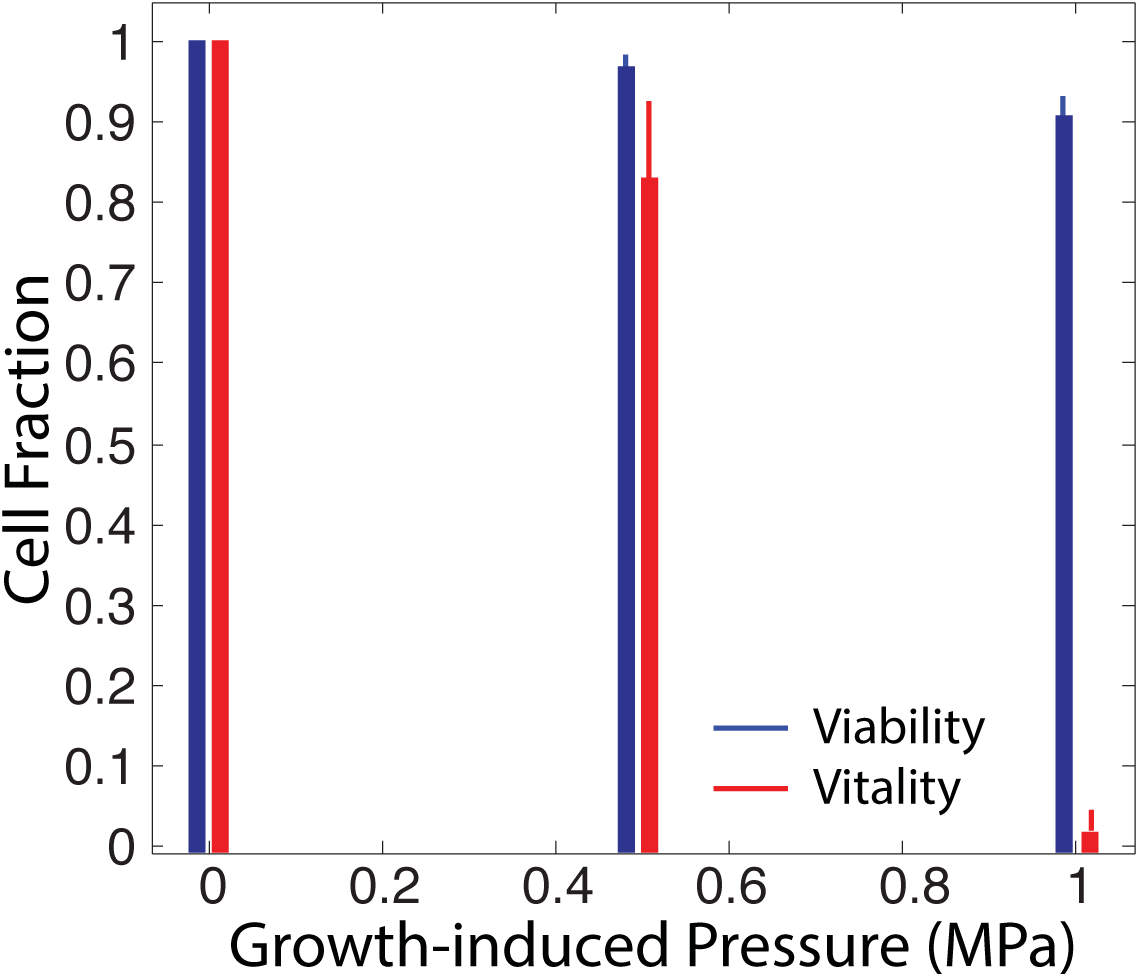
Measurement of cell viability and cell vitality. We assess how pressure changes cell viability and metabolic activity. Cell viability is assessed through a viability kit (LIVE/DEAD Yeast Viability Kit, Thermo Fisher Scientific). Briefly, propidium iodide (PI) is added to the culture medium. PI only enters the nucleus of dead cells and binds to DNA. We observe that, even at maximum pressure, most of the cells are alive (more than 90% of the cells). Cell vitality is assessed by flowing in a cell permeable esterase substrate (FungaLight Yeast CFDA, AM, Thermo Fisher Scientific) that is cleaved by esterases. The cleaved molecule becomes fluorescent, which enables one to assess esterase activity, which is directly linked to the global cell metabolic activity. We observe that, even though cell vitality does not change much at 0.5 MPa (the change is less than 15%), it is almost non existent at the maximum pressure of 1 MPa. This suggests that, even though alive, cells are not metabolically active. This could be explained by pressure-induced molecular crowding, as in [20], where all processes in the cell are slowed down to the point of stalling by the very high compression. Note that at the highest pressure, we observe about 5% of the cells bursting.

**Figure S5.**
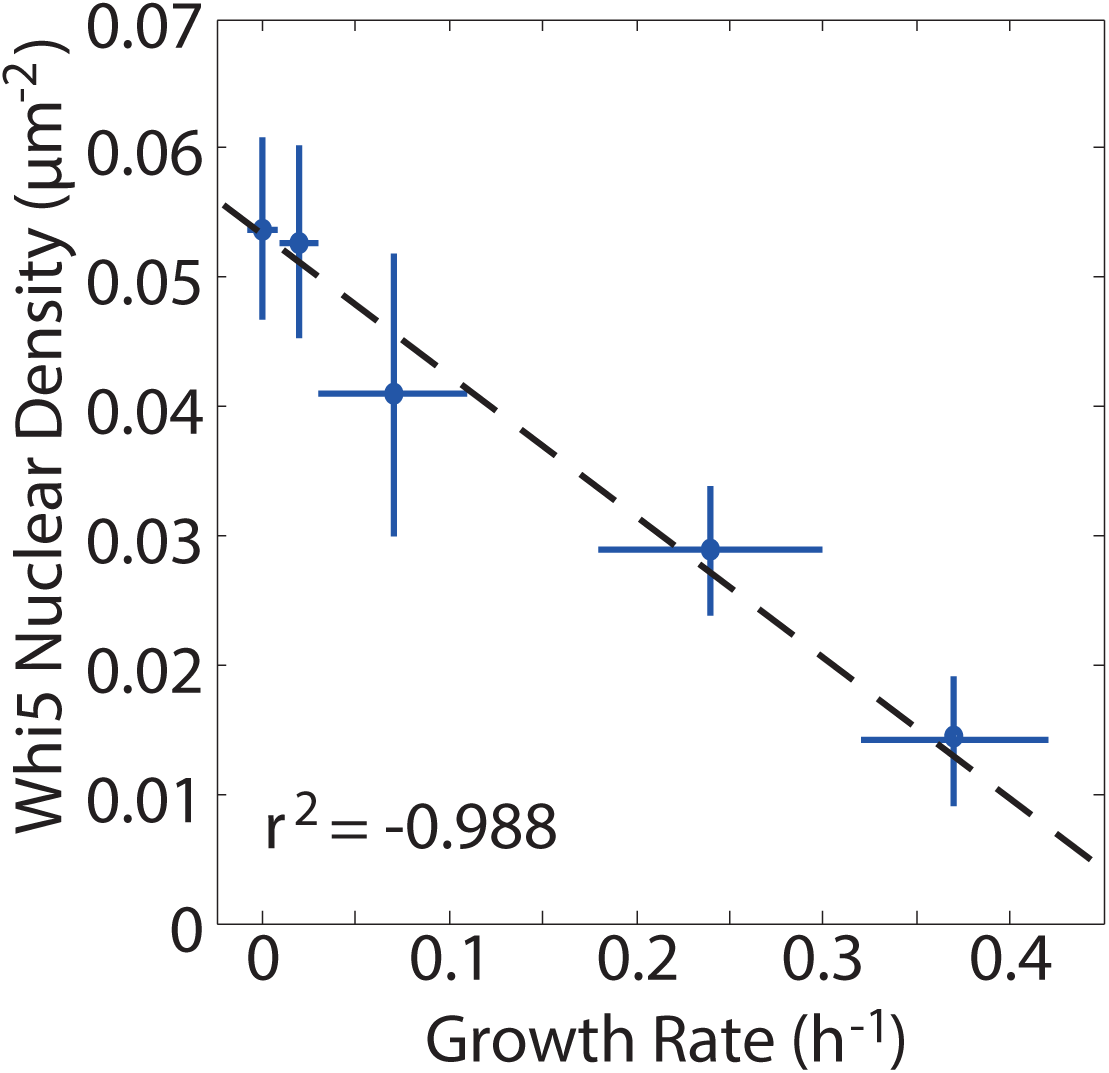
The density of nuclear Whi5 is anti-correlated with the growth rate. This plot shows the nuclear Whi5 density for different growth-induced pressures. The Whi5 density was obtained by measuring the number of cells presenting a nuclear Whi5 normalized by the observed area. Note that the nuclear density of Whi5 is increasing with decreasing growth rate, suggesting that growth rate reduction is accompanied with a cell cycle delay in the G1 phase of the cell cycle.

**Figure S6.**
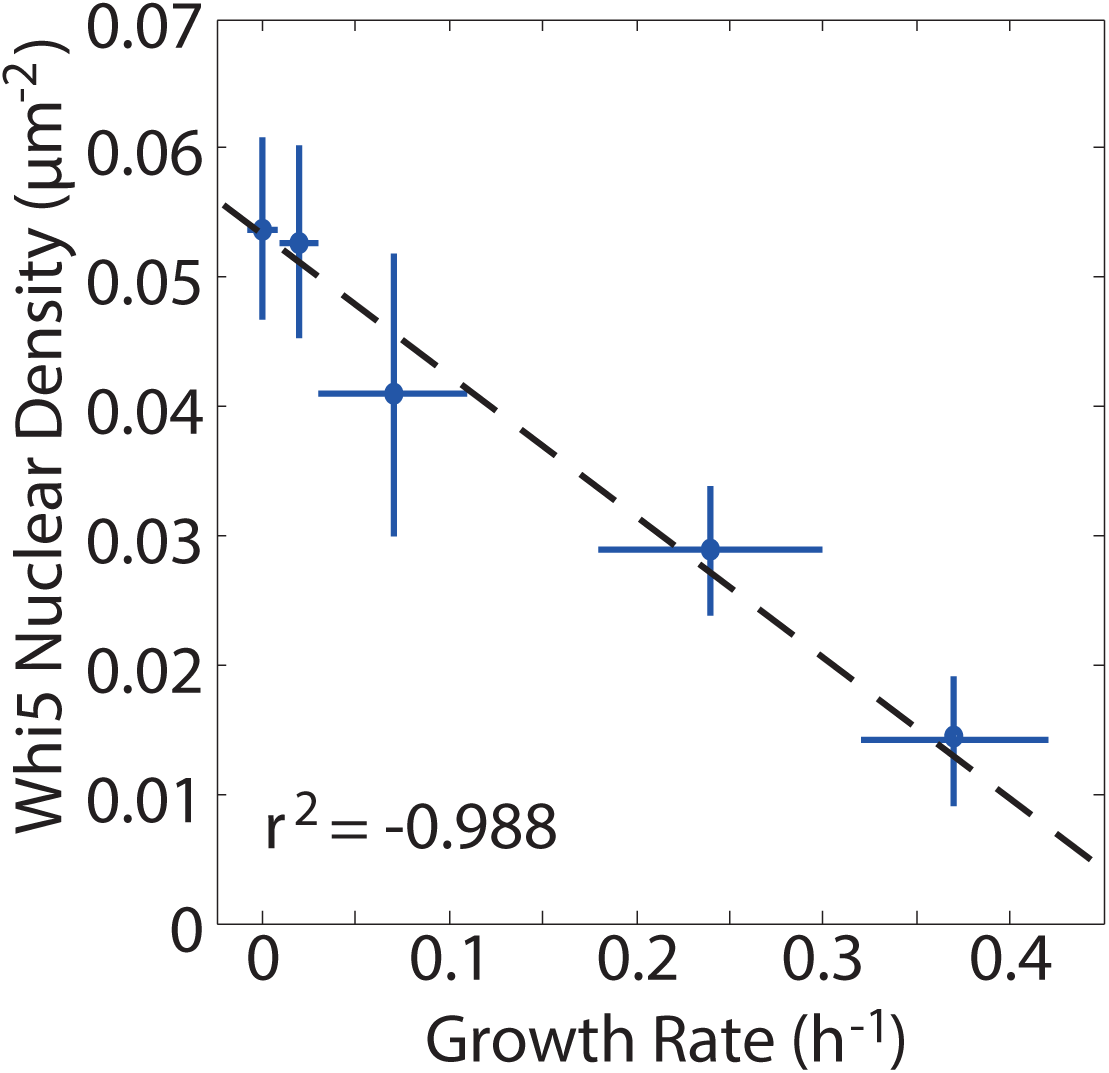
Relationship between fraction of surface covered and growth-induced pressure indicates turgor pressure adaptation. **A**. The growth-induced pressure increases (circles) faster than linearly with the fraction of surface covered. The dashed lines are obtained from our bead-spring simulations, in which yeast cells are modeled as elastic shells subject to a *constant* turgor pressure. The simulations yield a growth-induced pressure that increases linearly with surface coverage. The slope is equal to the turgor pressure Π, for which we chose three different values (see inset). The discrepancy between data and simulations suggests that the turgor pressure increases with growth-induced pressure. **B**. The growth-induced pressure divided by the fraction of covered surface corresponds to the pressure exerted in the contacts between cells and cover slip. Accordingly, the constant turgor pressure simulations of elastic shells yield nearly horizontal lines. The data, however, clearly shows that the pressure in the cell-coverslip contacts increase with the growth-induced pressure. This may indicate a gradual increase in turgor pressure. Error bars of the simulation data are smaller than the symbols. Error bars for the surface coverage are estimated as followed: We assume that we cannot measure the contact better than the diffraction limit. Hence, assuming a circular contact, we write that the radius of the contact has a typical error of ±*δ*, where *δ* is the radius of the Point Spread Function.

**Figure S7.**
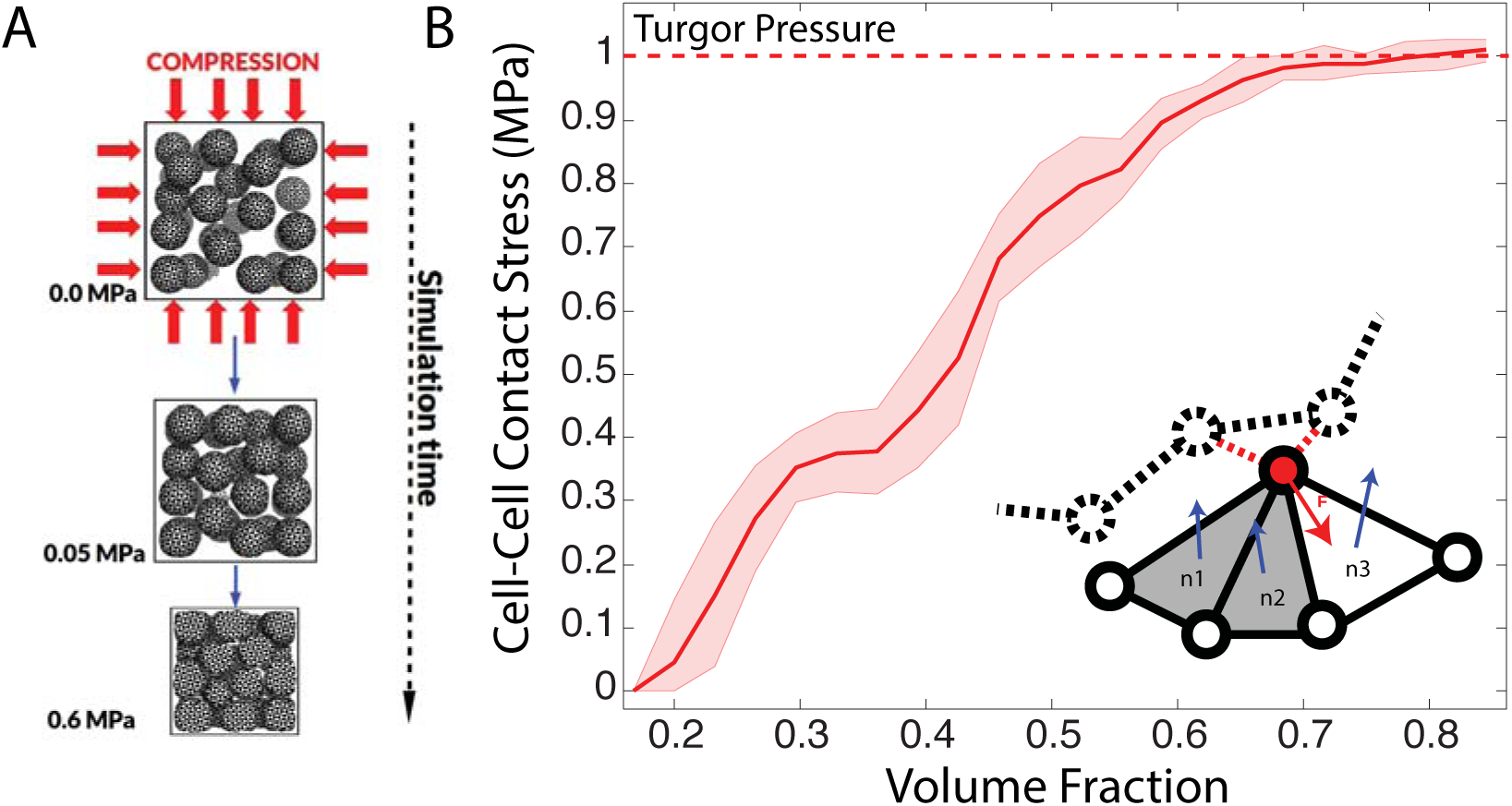
The average cell-cell contact stress approaches the cell turgor pressure under high compressive stress. We measure the cell-cell contact stress in mass-spring simulations, and find that for high compression / packing fraction, the stress approaches the internal cell turgor pressure. **A**. Mass-spring simulations scheme. Identical cells are randomly distributed in a rigid box. The initial concentration is low so the cells do not touch one another. Compression is performed by successive reduction of the size of the simulation box with the rate in the range of 0.01 - 0.1*μ*m*s*^−1^. **B**. 50 identical cells (*R*_0_ = 2.5*μ*m, Π = 1.0 MPa, E = 150 MPa, t = 0.1*μ*m) are compressed. For each pair of cells, the contact stress is calculated and the average contact stress is plotted (red line) versus the fraction of box volume occupied by cells. For strong compression (>0.7) the value of the average contact stress saturates at the value equal to turgor pressure, 1 MPa. The envelope corresponds to ±SD and is obtained out of 5 replicates simulations with different random initial cell positions and orientations. **Inset**. The contact stress σ_*c*_ is calculated as a ratio of the total normal force between two cells *F*_*n*_ and total contact area *A*_*c*_. The contact area *A*_*c*_ on one cell is a sum of areas of all triangles being in contact with the other cell. A triangle is in contact with another cell if all its vertices are in contact with the neighbor cell (non-zero repulsive forces). The total normal force exerted on one cell is a sum of all normal forces exerted on each vertex by the neighbor cell. To calculate the normal force **F** (red arrow) acting on a vertex (black-red circle), first the sum of all non-bonded repulsive forces, **F**_*rep*_ (red dashed lines), is calculated. Next the normal component of this force is extracted as a dot product with all the triangles (described by the normal vectors n;) being in contact with the neighboring cell (shaded triangles), **n***i* · **F**_*rep*_. In order to avoid double counting of the normal component of the force **F**_*rep*_ each dot product with **n***i* is scaled by the area of the triangle on which the force **F**_*rep*_ is acting.

**Figure S8.**
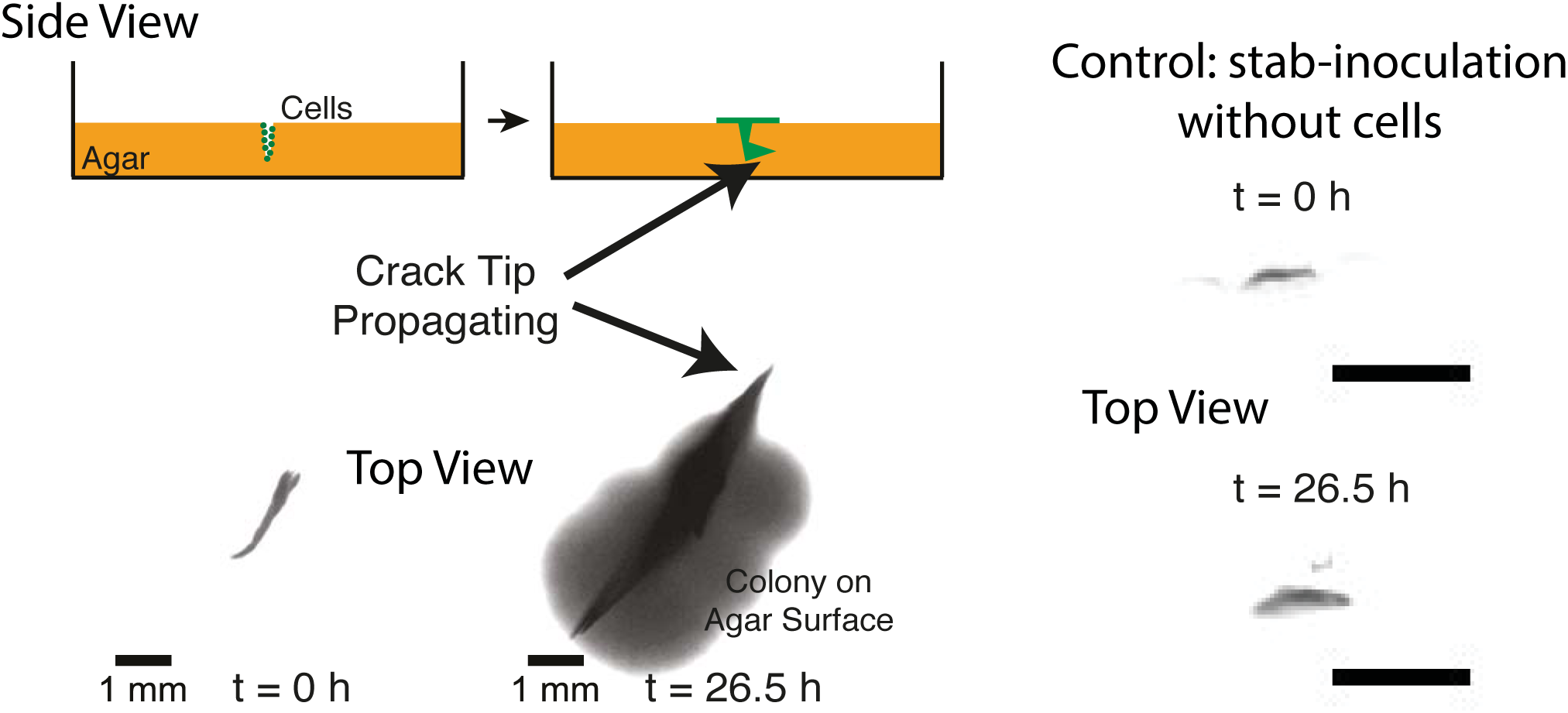
Self-driven jamming can propagate cracks in agar gels. We inoculate an agar gel (2%) by plunging in it a 0.45 mm diameter needle, which was first dipped in an overnight culture of budding yeast (strain S288C). The agar dish is then incubated at 30 degree Celsius under humidity control (to avoid drying). As the cartoon illustrates, cells flow out of the crack, and grow on the surface of the agar gel. The cells on top of the dish give rise to the large cloud on the lower image observed at 26.5 h, showing that the cells are not fully trapped in the crack. Nevertheless, the crack tips are propagating as a function of time, presumably due to jamming. As a control, we show images of cracks that were created by stabbing without cells and then incubated for the same amount of time. A time-lapse movie of the crack propagation is available **here**.

**Figure S9.**
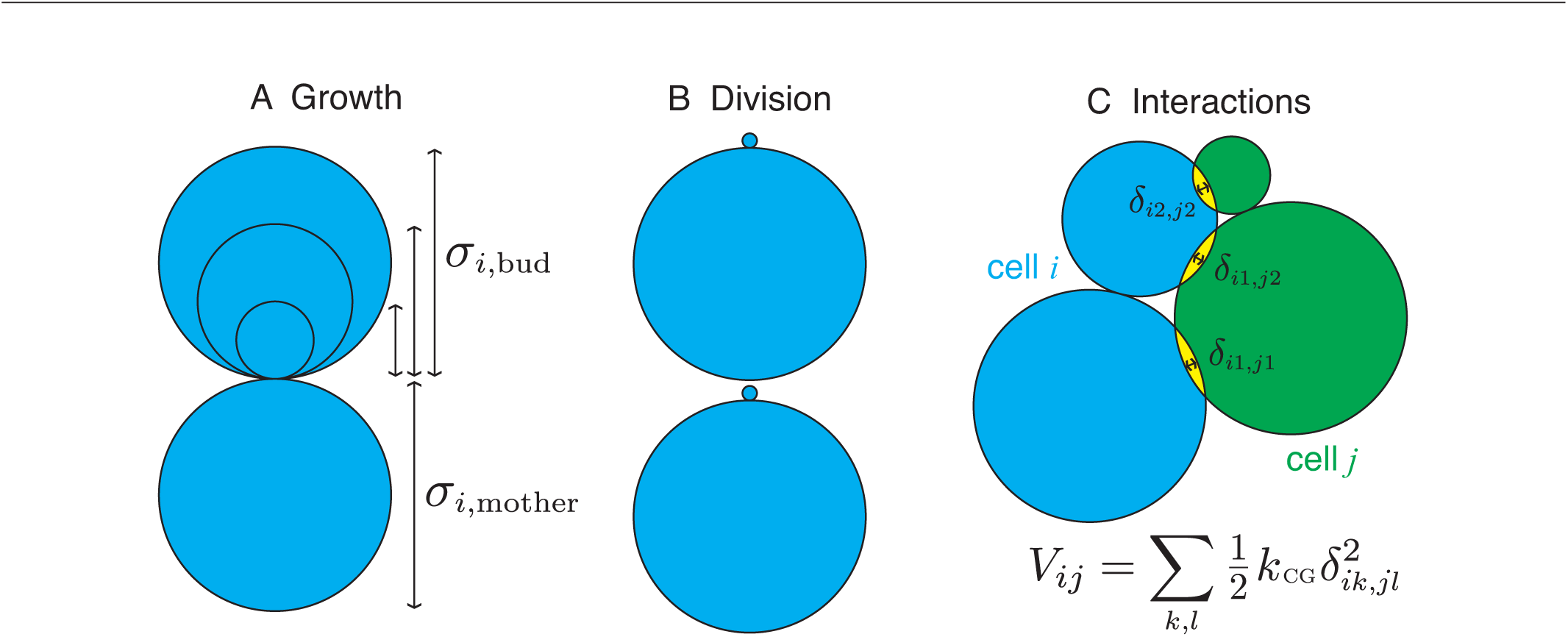
How cells grow in our coarse-grain simulation. Schematic of **A** the growth and **B** division processes and **C** inter-cell interactions in our coarse-grained simulations. Each cell is composed to two lobes, the mother and bud. **A**. During growth, the mother lobe diameter of cell *i* stays fixed at σ_*i*,mother_ = σ while the bud grows from σ_*i*,bud_ = 0 to σ_*i*,mother_ = σ. **B**. Once the bud reaches σ_*i*,mother_ = σ, cell *i* divides into two new daughter cells that retain the orientation of their mother cell. **C**. Cells *i* and *j* interact via only upon overlap via repulsive linear spring interactions with modulus *k*.

**Figure S10.**
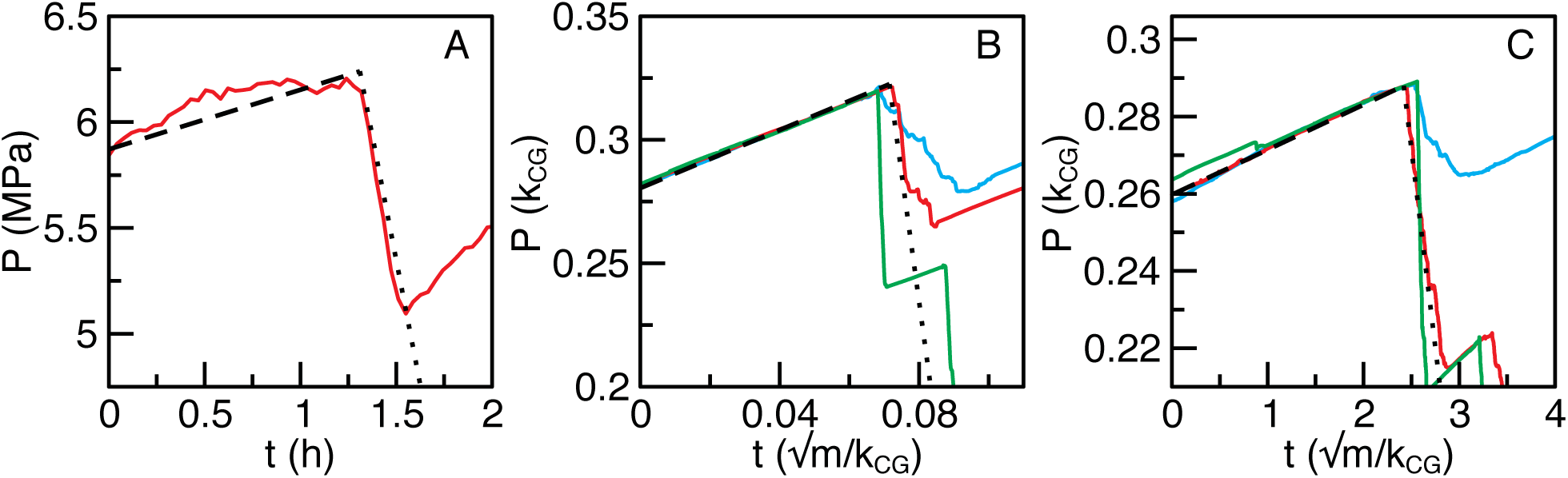
How we parameterize our corse-grain simulation. We use the experimental pressure curves to parameterize our coarse-grain simulations: the pressure rise enables to parameterize the growth, and the pressure drop the damping rate. Pressure as function of time during a single pressure drop for **A** experiments, **B** simulations without feedback, and **C** simulations with feedback (*P*_0_/*k* = 0.07). The red line in **A** corresponds to experiments with a 135° value. The red lines in **B** and **C** corresponds to simulations with best-fit values of *μ* 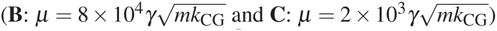 used in Fig. 2B and Fig. 2C of the main text, the cyan line corresponds to larger values of *μ* 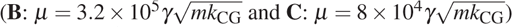 and the green line corresponds to smaller values of *μ* 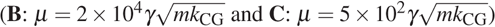. For **A**, the dashed line shows the mean slope during pressure increase 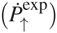 and the dotted line shows mean slope during avalanche 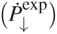. For **B** and **C**, the dashed line is the mean slope during increase 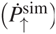 and the dotted line shows the extracted value of 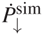 that yields the same ratio of *Ṗ*_↓_/*Ṗ*_↑_ as experiments.

## Footnotes

* Note that, because water can flow in and out of cells, hydrostatic pressures are conceptually very different from contact pressures studied here.

## References

[1] Paula Watnick and Roberto Kolter. Biofilm, city of microbes. Journal of bacteriology, 182(10):2675–2679, 2000.

[2] Saranna Fanning and Aaron P Mitchell. Fungal biofilms. PLoS Pathog, 8(4):e1002585, 2012.

[3] Vigdis Torsvik and Lise Øvreås. Microbial diversity and function in soil: from genes to ecosystems. Current opinion in microbiology, 5(3):240–245, 2002.

[4] Lionel Ranjard and Agnès Richaume. Quantitative and qualitative microscale distribution of bacteria in soil. Research in Microbiology, 152(8):707–716, 2001.

[5] James N Wilking, Vasily Zaburdaev, Michael De Voider, Richard Losick, Michael P Brenner, and David A Weitz. Liquid transport facilitated by channels in bacillus sub-tilis biofilms. Proceedings of the National Academy of Sciences, 110(3):848–852, 2013.

[6] James N Wilking, Thomas E Angelini, Agnese Seminara, Michael P Brenner, and David A Weitz. Biofilms as complex fluids. MRS bulletin, 36(05):385–391, 2011.

[7] Munehiro Asally, Mark Kittisopikul, Pau Rué, Yingjie Du, Zhenxing Hu, Tolga Çaǧatay, Andra B Robinson, Hongbing Lu, Jordi Garcia-Ojalvo, and Gürol M Süel. Localized cell death focuses mechanical forces during 3d patterning in a biofilm. Proceedings of the National Academy of Sciences, 109(46): 18891–18896, 2012.

[8] Alfred B Cunningham, William G Characklis, Feisal Abe-deen, and David Crawford. Influence of biofilm accumulation on porous media hydrodynamics. Environmental science & technology, 25(7): 1305–1311, 1991.

[9] Bruce E Rittmann. The significance of biofilms in porous media. Water Resources Research, 29(7):2195–2202, 1993.

[10] Neil AR Gow, Alistair JP Brown, and Frank C Odds. Fungal morphogenesis and host invasion. Current opinion in microbiology, 5(4):366–371, 2002.

[11] Nicholas P Money. Turgor pressure and the mechanics of fungal penetration. Canadian journal of botany, 73(S1):96–102, 1995.

[12] Timothy J Foster, Joan A Geoghegan, Vannakambadi K Ganesh, and Magnus Höök. Adhesion, invasion and evasion: the many functions of the surface proteins of staphylococcus aureus. Nature Reviews Microbiology, 12(1):49–62, 2014.

[13] Alexander E Smith, Zhibing Zhang, Colin R Thomas, Kennith E Moxham, and Anton PJ Middelberg. The mechanical properties of saccharomyces cerevisiae. Proceedings of the National Academy of Sciences, 97(18):9871–9874, 2000.

[14] Nicolas Mine, Arezki Boudaoud, and Fred Chang. Mechanical forces of fission yeast growth. Current Biology, 19(13): 1096–1101, 2009.

[15] JD Stenson, CR Thomas, and P Hartley. Modelling the mechanical properties of yeast cells. Chemical Engineering Science, 64(8): 1892–1903, 2009.

[16] Hannah H Tuson, George K Auer, Lars D Renner, Mariko Hasebe, Carolina Tropini, Max Salick, Wendy C Crone, Ajay Gopinathan, Kerwyn Casey Huang, and Douglas B Weibel. Measuring the stiffness of bacterial cells from growth rates in hydrogels of tunable elasticity. Molecular microbiology, 84(5):874–891, 2012.

[17] Farhang Radjai, Michel Jean, Jean-Jacques Moreau, and Stéphane Roux. Force distributions in dense two-dimensional granular systems. Physical review letters, 77(2):274, 1996.

[18] Trushant S Majmudar and Robert P Behringer. Contact force measurements and stress-induced anisotropy in granular materials. Nature, 435(7045): 1079–1082, 2005.

[19] Dapeng Bi, Jie Zhang, Bulbul Chakraborty, and RP Behringer. Jamming by shear. Nature, 480(7377):355–358, 2011.

[20] Claus Heussinger and Jean-Louis Barrat. Jamming transition as probed by quasistatic shear flow. Physical review letters, 102(21):218303, 2009.

[21] Th Warscheid and Joanna Braams. Biodeterioration of stone: a review. International Biodeterioration & Biodegradation, 46(4):343–368, 2000.

[22] Claus Seebacher, Jochen Brasch, Dietrich Abeck, Oliver Comely, Isaak Effendy, Gabriele Ginter-Hanselmayer, Norbert Haake, Gudrun Hamm, U-Ch Hipler, Herbert Hof, et al. Onychomycosis. Mycoses, 50(4):321–327, 2007.

[23] L Julia Douglas. Candida biofilms and their role in infection. Trends in microbiology, 11(1):30–36, 2003.

[24] GDR MiDi. On dense granular flows. The European Physical Journal E, 14(4):341–365, 2004.

[25] Luanne Hall-Stoodley, J William Costerton, and Paul Stoodley. Bacterial biofilms: from the natural environment to infectious diseases. Nature Reviews Microbiology, 2(2):95–108, 2004.

[26] Jin-Ah Park, Jae Hun Kim, Dapeng Bi, Jennifer A Mitchel, Nader Taheri Qazvini, Kelan Tantisira, Chan Young Park, Maureen McGill, Sae-Hoon Kim, Bomi Gweon, et al. Unjamming and cell shape in the asthmatic airway epithelium. Nature materials, 2015.

[27] Amy Rowat, James Bird, Jeremy Agresti, Oliver Rando, and David Weitz. Tracking lineages of single cells in lines using a microfluidic device. Proceedings of the National Academy of Sciences, 106(43): 18149–18154, 2009.

[28] H Cho, Henrik Jönsson, Kyle Campbell, Pontus Melke, Joshua W Williams, Bruno Jedynak, Ann M Stevens, Alex Groisman, and Andre Levchenko. Self-organization in high-density bacterial colonies: efficient crowd control. PLoS Biol, 5(11):e302, 2007.

[29] Frederick K Balagaddé, Lingchong You, Carl L Hansen, Frances H Arnold, and Stephen R Quake. Long-term monitoring of bacteria undergoing programmed population control in a microchemostat. Science, 309(5731): 137–140, 2005.

[30] Gilles Charvin, Frederick R Cross, Eric D Siggia, et al. A microfluidic device for temporally controlled gene expression and long-term fluorescent imaging in unperturbed dividing yeast cells. Plos one, 3(1):e1468–e1468, 2008.

[31] IM De Mara on, Pierre-André Marechal, and Patrick Gervais. Passive response of saccharomyces cerevisiae to osmotic shifts: cell volume variations depending on the physiological state. Biochemical and biophysical research communications, 227:519–523, 1996.

[32] Iker Zuriguel, Angel Garcimartín, Diego Maza, Luis A Pugnaloni, and JM Pastor. Jamming during the discharge of granular matter from a silo. Physical Review E, 71(5):051303, 2005.

[33] M. E. Cates, J. P. Wittmer, Bouchaud J.-P, and P. Claudin. Jamming, force chains, and fragile matter. Physical review letters, 81(9):1841, 1998.

[34] Markus Basan, Thomas Risler, Jean-François Joanny, Xavier Sastre-Garau, and Jacques Prost. Homeostatic competition drives tumor growth and metastasis nucle-ation. HFSP journal, 3(4):265–272, 2009.

[35] Morgan Delarue, Fabien Montel, Danijela Vignjevic, Jacques Prost, Jean-François Joanny, and Giovanni Cap-pello. Compressive stress inhibits proliferation in tumor spheroids through a volume limitation. Biophysical journal, 107(8): 1821–1828, 2014.

[36] D Weaire and MA Fortes. Stress and strain in liquid and solid foams. Advances in Physics, 43(6):685–738, 1994.

[37] Yves F Dufrêne. Sticky microbes: forces in microbial cell adhesion. Trends in microbiology, 2015.

[38] David R Soil. Candida biofilms: is adhesion sexy? Current Biology, 18(16):R717–R720, 2008.

[39] Libuše Váchová, Vratislav Št’ovíček, Otakar Hlaváček, Oleksandr Chernyavskiy, Luděk Štěpánek, Lucie Kubínová, and Zdena Palková. Flollp, drug efflux pumps, and the extracellular matrix cooperate to form biofilm yeast colonies. The Journal of cell biology, 194(5):679–687, 2011.

[40] Adem Dogangun, Zeki Karaca, Ahmet Durmus, and Halil Sezen. Cause of damage and failures in silo structures. Journal of performance of constructed facilities, 23(2):65–71, 2009.

## References

[1] D Brambley, B Martin, and PD Prewett. Microlithography: an overview. Advanced Materials for Optics and Electronics, 4(2):55–74, 1994.

[2] Zhigang Wu and Klas Hjort. Surface modification of pdms by gradient-induced migration of embedded pluronic. Lab on a Chip, 9(11): 1500–1503, 2009.

[3] Robert A Preston, Robert F Murphy, and Elizabeth W Jones. Apparent endocytosis of fluorescein isothiocyanate-conjugated dextran by saccharomyces cerevisiae reflects uptake of low molecular weight impurities, not dextran. The Journal of cell biology, 105(5): 1981–1987, 1987.

[4] IM De Mara on, Pierre-André Marechal, and Patrick Gervais. Passive response of saccharomyces cerevisiae to osmotic shifts: cell volume variations depending on the physiological state. Biochemical and biophysical research communications, 227:519–523, 1996.

[5] JD Stenson, CR Thomas, and P Hartley. Modelling the mechanical properties of yeast cells. Chemical Engineering Science, 64(8): 1892–1903, 2009.

[6] John D Stenson, Peter Hartley, Changxiang Wang, and Colin R Thomas. Determining the mechanical properties of yeast cell walls. Biotechnology progress, 27(2):505–512, 2011.

[7] WW Feng and W-H Yang. On the contact problem of an inflated spherical nonlinear membrane. Journal of Applied Mechanics, 40(1):209–214, 1973.

[8] Alexander E Smith, Zhibing Zhang, Colin R Thomas, Kennith E Moxham, and Anton PJ Middelberg. The mechanical properties of saccharomyces cerevisiae. Proceedings of the National Academy of Sciences, 97(18):9871–9874, 2000.

[9] Maciej Kot, Hiroshi Nagahashi, and Piotr Szymczak. Elastic moduli of simple mass spring models. The Visual Computer, pages 1–12, 2014.

[10] Carl F Schreck, Ning Xu, and Corey S O’Hern. A comparison of jamming behavior in systems composed of dimer-and ellipse-shaped particles. Soft Matter, 6(13):2960–2969, 2010.

[11] FDC Farrell, O Hallatschek, D Marenduzzo, and B Waclaw. Mechanically driven growth of quasi-two-dimensional microbial colonies. Physical review letters, 111(16):168101, 2013.

[12] Dmitri Volfson, Scott Cookson, Jeff Hasty, and Lev S Tsimring. Biomechanical ordering of dense cell populations. Proceedings of the National Academy of Sciences, 105(40):15346–15351, 2008.

[13] Ehsan Irani, Pinaki Chaudhuri, and Claus Heussinger. Impact of attractive interactions on the rheology of dense athermal particles. Physical review letters, 112(18):188303, 2014.

[14] Gregg Lois, Jerzy Blawzdziewicz, and Corey S OHern. Jamming transition and new percolation universality classes in particulate systems with attraction. Physical review letters, 100(2):028001, 2008.

[15] Karine Guevorkian, Marie-Josée Colbert, Mélanie Durth, Sylvie Dufour, and Françoise Brochard-Wyart. Aspiration of biological viscoelastic drops. Physical review letters, 104(21):218101, 2010.

[16] Céline Bottier, Chiara Gabella, Benoî Vianay, Lara Buscemi, Ivo F Sbalzarini, Jean-Jacques Meister, and Alexander B Verkhovsky. Dynamic measurement of the height and volume of migrating cells by a novel fluorescence microscopy technique. Lab on a Chip, 11(22):3855–3863, 2011.

[17] Brian S Hardy, Kawika Uechi, Janet Zhen, and H Pirouz Kavehpour. The deformation of flexible pdms microchannels under a pressure driven flow. Lab on a Chip, 9(7):935–938, 2009.

[18] N Ertugay, H Hamamci, and A Bayindirli. Fed-batch cultivation of bakers yeast: effect of nutrient depletion and heat stress on cell composition. Folia microbiologica, 42(3):214–218, 1997.

[19] Hyun Youk and Alexander van Oudenaarden. Growth landscape formed by perception and import of glucose in yeast. Nature, 462(7275):875–879, 2009.

[20] Pascal Hersen, Megan N McClean, L Mahadevan, and Sharad Ramanathan. Signal processing by the hog map kinase pathway. Proceedings of the National Academy of Sciences, 105(20):7165–7170, 2008.

